# Evaluation of hybrid and non-hybrid methods for de novo assembly of nanopore reads

**DOI:** 10.1101/030437

**Authors:** Ivan Sović, Krešimir Križanović, Karolj Skala, Mile Šikić

## Abstract

Recent emergence of nanopore sequencing technology set a challenge for the established assembly methods not optimized for the combination of read lengths and high error rates of nanopore reads. In this work we assessed how existing *de novo* assembly methods perform on these reads. We benchmarked three non-hybrid (in terms of both error correction and scaffolding) assembly pipelines as well as two hybrid assemblers which use third generation sequencing data to scaffold Illumina assemblies. Tests were performed on several publicly available MinION and Illumina datasets of *E. coli* K-12, using several sequencing coverages of nanopore data (20x, 30x, 40x and 50x). We attempted to assess the quality of assembly at each of these coverages, to estimate the requirements for closed bacterial genome assembly. Results show that hybrid methods are highly dependent on the quality of NGS data, but much less on the quality and coverage of nanopore data and perform relatively well on lower nanopore coverages. Furthermore, when coverage is above 40x, all non-hybrid methods correctly assemble the *E. coli* genome, even a non-hybrid method tailored for Pacific Bioscience reads. While it requires higher coverage compared to a method designed particularly for nanopore reads, its running time is significantly lower.

## 1 Introduction

During the last ten years Next generation sequencing (NGS) devices have dominated genome sequencing market. In contrast to previously used Sanger sequencing, NGS is much cheaper, less time consuming and not so labour intensive. Yet, when it comes to de novo assembly of longer genomes many researchers are being sceptical of using NGS reads. These devices produce reads a few hundred base pairs long, which is too short to unambiguously resolve repetitive regions even within relatively small microbial genomes (Nagarajan and Pop, 2013).

Although usage of paired-end and mate-pair technologies has improved the accuracy and completeness of assembled genomes, NGS sequencing still produces highly fragmented assemblies and is currently mostly employed for deep resequencing. Nonetheless, owing to NGS many efficient algorithms have been developed to optimize the running time and memory footprints in sequence assembly, alignment and downstream analysis steps.

The need for the technologies that would produce longer reads which could solve the problem of repeating regions has resulted in the advent of new sequencing approaches – the so-called “third generation sequencing technologies”. The first among them was single-molecule sequencing technology developed by Pacific Biosciences (PacBio). Although PacBio sequencers produce much longer reads (up to several tens of thousands of base pairs), their reads have a significantly higher error (~10-15%) rate than NGS reads (<2%) (Nagarajan and Pop, 2013). Existing assembly and alignment algorithms were not capable of handling such high error rates. This caused the development of read error correction methods. At first hybrid correction was performed using complementary NGS (Illumina) data (Koren *et al*., 2012). Later, self-correction of PacBio-only reads was developed (Chin *et al*., 2013) which requires higher coverage (>50x). The development of new, more sensitive aligners was required (BLASR (Chaisson and Tesler, 2012) and optimization of existing ones (BWA-MEM (Li, 2013)).

In 2014, Oxford Nanopore Technologies (ONT) presented their tiny MinION sequencer - about the size of a harmonica. The MinION can produce reads up to a few hundred thousand base pairs long, but with a higher error rate compared to PacBio reads. For example, 1D reads from the MinION sequencer have raw base accuracy less than 65-75%; higher quality 2D reads (80-88% accuracy) comprise a fraction of all 2D reads and even smaller fraction of the total dataset, with overall median accuracy being between 70-85% (Ashton *et al*., 2014; Laver *et al*., 2015; Ip *et al*., 2015). This again spurred the need for development of even more sensitive algorithms for mapping such as GraphMap (Sovic *et al*., 2015) and realignment marginAlign (Jain *et al*., 2015). Any doubt about the possibility of using MinION reads for de novo assembly was resolved in 2015 when Loman et al. demonstrated (Loman *et al*., 2015) that the assembly of a bacterial genome (E. Coli K-12) using solely ONT reads is possible in spite of high error rates. Thanks to the extremely long reads and the affordability and availability of the nanopore sequencing technology, these results might cause a revolution in de novo sequence analysis.

Following up on the results from Loman et al. (Loman *et al*., 2015) and Liao et al (Liao *et al*., 2015), we explored the applicability of existing hybrid and non-hybrid de novo assembly tools that support third generation sequencing data and assessed their ability to cope with nanopore error profiles. In our study, we compared five assembly tools/pipelines which include three long-read assemblers: pipeline published by Loman et al. (in continuation LQS pipeline), PBcR (Koren *et al*., 2012), Falcon; and two hybrid assemblers: ALLPATHS-LG (Gnerre *et al*., 2011) and SPAdes (Bankevich *et al*., 2012). These tools were tested on real, publicly available datasets of a well-known clonal sample of *E. coli* K-12 MG1655. All of the tools/pipelines were tested up to the draft assembly level, not including the polishing phase.

## 2 Background

Majority of algorithms for de novo assembly follow either the de Bruijn graph (DBG) or the Overlap-Layout-Consensus (OLC) paradigm (Pop, 2009). While the early assemblers, specialized for Sanger sequencing methods - such as Celera - use the OLC paradigm, de novo assemblers developed for NGS data are mostly based on the DBG approach with several exceptions. Although the DBG approach is faster, OLC based algorithms perform better for longer reads (Pop, 2009),. Additionally, the DBG assemblers depend on finding exact-matching kmers between reads (typically ~20-63 bases long). Given the error rates in the third generation sequencing data, this presents a serious limitation. On the other hand, the OLC approach should be able to cope with higher error rates given a sensitive enough overlapper, but contrary to DBG the time-consuming all-to-all pairwise comparison between the input reads still needs to be performed.

Since the focus in the past decade has been on the NGS reads, there are not many OLC assemblers that could be utilized for long PacBio and ONT reads. In fact, the methods developed to handle such data are mostly pipelines based on the Celera assembler, including: HGAP (Chin *et al*., 2013), PBcR (Koren *et al*., 2012) and the LQS pipeline (Loman *et al*., 2015). Since its original publication (Myers *et al*., 2000), Celera has been heavily revised to support newer sequencing technologies, including modifications for second generation data (Miller *et al*., 2008), adoptions for the third generation (single molecule) data via hybrid error correction (Koren *et al*., 2012), non-hybrid error correction (Miller *et al*., 2008; Berlin *et al*., 2015) and hybrid approaches to assembly which combine two or more technologies (Goldberg *et al*., 2006). Celera was used in numerous sequencing projects, including the Drosophila assembly and the assembly of both parental haplotypes from a single human genome (Levy *et al*., 2007). All of this contributed to the popularity of Celera which led to its wide adoption in pipelines for the assembly of third generation sequencing data. Notably, one of the first was the Hierarchical Genome Assembly Process (HGAP) which is not only an assembly pipeline but also a specification of a workflow which can be implemented using various components at each step. There are currently three HGAP implementations (Pacific Biosciences, 2013): HGAP in SMRT Analysis, HBAR-DTK development toolkit and PacBioToCA implemented in the WGS (Celera) package. HGAP uses BLASR to detect overlaps between raw reads during the error correction step. However, HGAP requires input data to be in the PacBio-specific formats, which limits its application to other (e.g. nanopore) sequencing technologies. PBcR, since recently, employs the MHAP overlapper (Berlin *et al*., 2015) for sensitive overlapping of reads during the error-correction step. Also, recent updates to PBcR allow it to handle reads from Oxford Nanopore MinION sequencers. The LQS pipeline follows a similar workflow to that of HGAP, but with novel error-correction (Nanocorrect) and consensus (Nanopolish) steps. Instead of BLASR and MHAP, Nanocorrect uses DALIGNER (Myers, 2014) for overlap detection. Nanopolish presents a new signal-level consensus method for fine-polishing of the draft assembly using raw nanopore data. The LQS pipeline also employs Celera as the middle layer, i.e. for assembly of the error corrected reads.

The only non-hybrid alternative to the Celera-based pipelines is Falcon (Biosciences, 2015). Falcon is a new experimental diploid assembler developed by Pacific Biosciences, not yet officially published. It is based on a hierarchical approach similar to HGAP, consisting of several steps: (I) raw sub-read overlapping for error correction using DALIGNER, (II) pre-assembly and error correction, (III) overlapping of error-corrected reads, (IV) filtering of overlaps, (V) construction of the string graph and (VI) contig construction. Unlike HGAP, it does not use Celera as its core assembler. Since Falcon accepts input reads in the standard FASTA format (and not only the PacBio-specific format like HGAP), it can potentially be used on any bascalled long-read dataset. Although originally intended for PacBio data, Falcon presents a viable option for assembly of nanopore reads, which exhibit slightly higher error rates than PacBio, but have notably different error profiles.

Aside from Celera-based assembly pipelines and Falcon, hybrid assembly approaches present another avenue to utilizing nanopore sequencing data. Liao et al. (Liao *et al*., 2015) recently evaluated several assembly tools on PacBio data, including hybrid assemblers SPAdes (Bankevich *et al*., 2012) and ALLPATHS-LG (Gnerre *et al*., 2011) for which they reported good results. Both of these use Illlumina libraries for the primary assembly, and then attempt to scaffold the assemblies using longer, less accurate reads. Furthermore, SPAdes was recently updated and now officially supports nanopore sequencing data as the long read complement to NGS data.

## 3 Methods

Since to the best of our knowledge no dedicated MinION read simulator exists, we focus our benchmark on real nanopore sequencing datasets. Although, there is a number of publicly available datasets, many of them consist either of organisms/strains which do not yet have officially finished genome assemblies, or the coverage of the dataset is not high enough to provide informative nanopore-only assembly results. Aside from the Lambda phage (which comes as a burn-in sample for every MinION), the most abundant are sequencing data for the well-known clonal sample of *E. coli* K-12 MG1655. In this study, we use several most recent *E. coli* K-12 datasets to reflect on the current state of the nanopore data as well as the quality of assembly they provide. In addition to using the entire datasets, we subsampled some of the datasets to provide a larger span of coverages in order to inspect the scalability of assemblers as well as their ability to cope with the abundance of the data.

### 3.1 Datasets

Benchmarking datasets were extracted from several publicly available nanopore datasets and one publicly available Illumina dataset. These include:

1. **ERX708228, ERX708229, ERX708230, ERX708231:** 4 flowcells used in Loman et al. nanopore assembly paper (Loman *et al*., 2015).
2. ***E. coli* K-12 MG1655 R7.3** dataset (Quick *et al*., 2014).
3. **MARC, WTCHG dataset** (Ip *et al*., 2015): A dataset recently published by the MinION Analysis and Reference Consortium, consists of a compilation of data generated using several MinION sequencers in laboratories distributed world-wide.
4. ***E. coli* K-12 MG1655 SQK-MAP006-1 dataset:** This is the most recent publicly available MinION dataset, obtained using the newest sequencing protocol. Link: http://lab.loman.net/2015/09/24/first-sqk-map-006-experiment/
5. **Illumina frag and jump libraries** (Liao *et al*., 2015): Link: ftp://ftp.broadinstitute.org/pub/papers/assembly/Ribeiro2012/data/ecoli_data_alt.tar.gz

The datasets were designed with the idea to test the effect of varying coverage and data quality on the assembly process, and consist of either full datasets described above or subsampled versions of these datasets.

Test dataset used for benchmarking include:

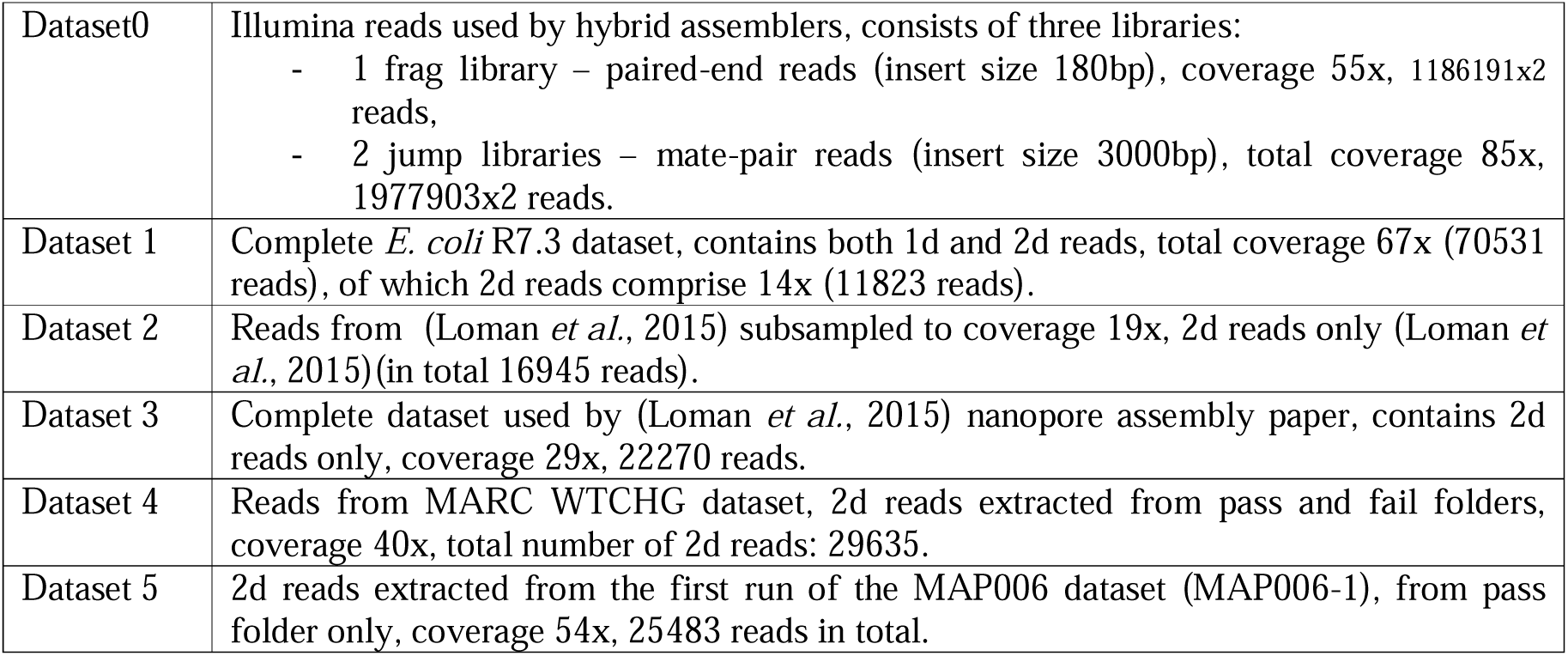

### 3.2 Data preparation

For nanopore datasets, sequence data was extracted from basecalled FAST5 files using Poretools (Loman and Quinlan, 2014). Data was extracted from a subset of folders or flowcells to achieve the desired coverage (e.g. for Dataset 2, flowcells ERX708228, ERX708229 and ERX708230 were used to obtain coverage close to 20x).

Hybrid assemblers were tested using the Illumina dataset together with each nanopore test dataset. They were also run on the Illumina dataset alone, to get a reference for assembly quality and to be able to estimate the contribution to assembly when nanopore reads are added. All libraries in the Illumina dataset come with reads and quality values in separate files (fasta and quala files). These were combined into fastq format using convertFastaAndQualToFastq.jar script downloaded from http://www.cbcb.umd.edu/software/PBcR/data/convertFastaAndQualToFastq.jar.

### 3.3 Assembly pipelines

**LQS pipeline**: Pipeline developed and published by Loman *et al*. in their pivotal nanopore assembly paper (Loman *et al*., 2015) (https://github.com/jts/nanopore-paper-analysis). The pipeline consists of Nanocorrect, WGS and Nanopolish. The version of the pipeline tested in this paper uses Nanocorrect commit 47dcd7f147c, WGS version 8.2 and Nanopolish commit 6440bfbfcf4fa.

**PBcR:** Implemented as a part of the WGS package (http://wgs-assembler.sourceforge.net/wiki/index.php/PBcR). In this paper version 8.3rc2 of WGS was used. Spec file defining assembly parameters for nanopore data, was downloaded from the PBcR web page.

**FALCON:** To evaluate Falcon we used the FALCON-integrate project (https://github.com/PacificBiosciences/FALCON-integrate) (commit: 3e7dd7db190). Since no formal parameter specification for nanopore data currently exists, we derived a suitable set of parameters through trial and error (**Suppl. Note 1**).

**SPAdes:** SPAdes v3.6.1 was downloaded from http://bioinf.spbau.ru/en/content/spades-download-0.

**ALLPATHS-LG:** ALLPATHS-LG release 52488 was downloaded from https://www.broadinstitute.org/software/allpaths-lg/blog/?page_id=12.

### 3.4 Evaluating the results

All assembly results were compared to the *E. coli* K-12 MG1655 NCBI reference, NC_000913.3. Assembly quality was evaluated using Quast 3.1 (Gurevich *et al*., 2013) and Dnadiff (Kurtz *et al*., 2004) tools. CPU and memory consumption was evaluated using a fork of the Cgmemtime tool (https://github.com/isovic/cgmemtime.git). For assemblies that produced one “big contig”, over 4Mpb in length, that contig was extracted and solely compared to the reference using Dnadiff tool.

## 4 Results

### 4.1 Assembly quality

Since Nanopolish currently does not support 1d reads, and Falcon and PBcR do not include a polishing phase, we focused on comparison of only the non-polished draft assemblies.

**Table 1** displays assembly results on datasets 2-5 assessed using Quast and Dnadiff tools. Dataset 1 analysis is omitted because of its particular characteristics. It has a greater total coverage but much lower data quality compared to other datasets (because of 1d reads). None of the non-hybrid assemblers managed to produce a good assembly using Dataset 1 (**Suppl. Table 1**), while both hybrid assemblers were able to use it to improve their assembly. It can be concluded that low 2d coverage together with high coverage of low quality 1d reads is not sufficient to complete an assembly of a bacterial genome using currently available methods.

**Table 1.**
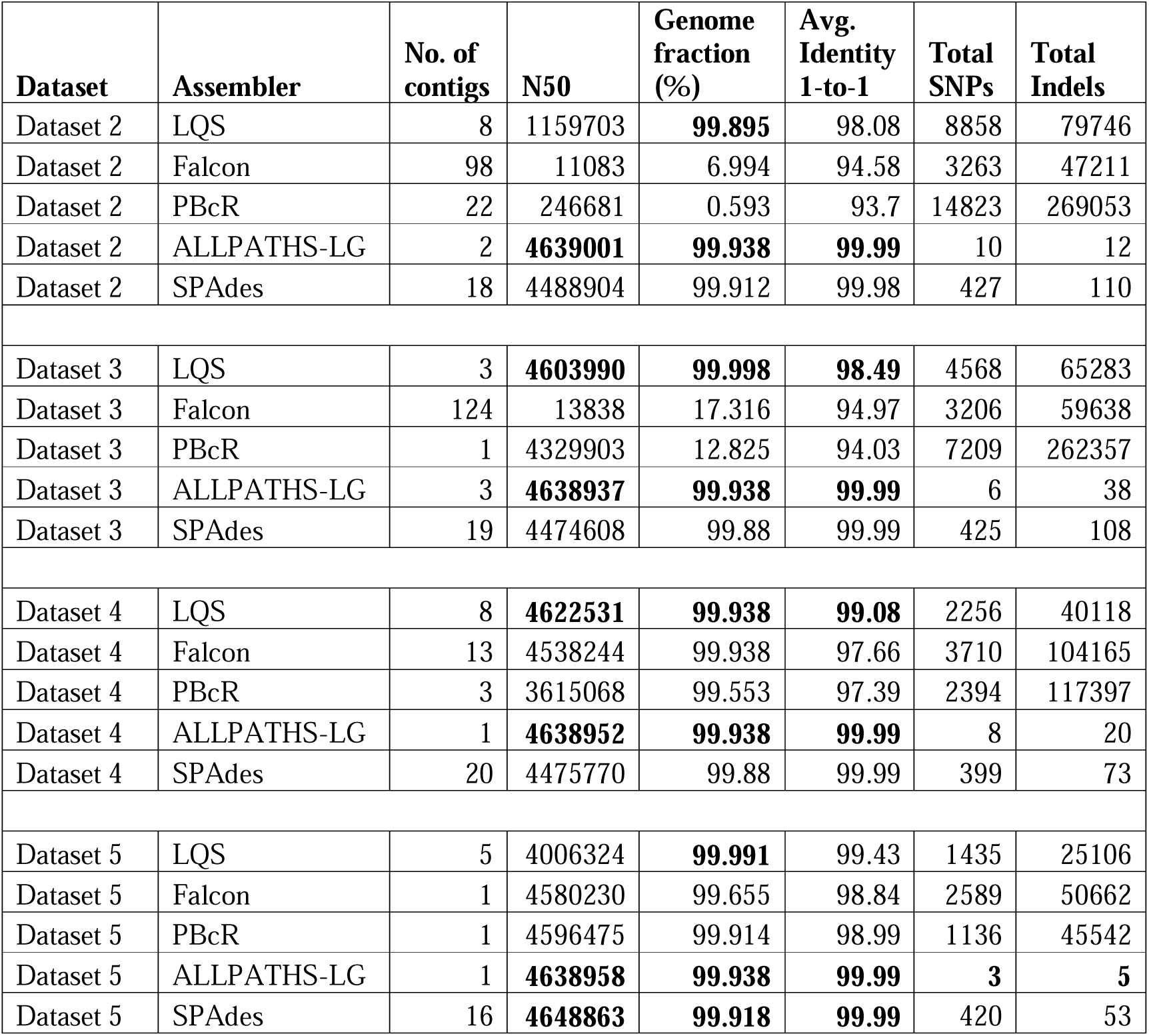
Assembly quality assessment using Quast and Dnadiff

| **Dataset** | **Assembler** | **No. of contigs** | **N50** | **Genome fraction (%)** | **Avg. Identity 1-to-1** | **Total SNPs** | **Total Indels** |
| --- | --- | --- | --- | --- | --- | --- | --- |
| Dataset 2 | LQS | 8 | 1159703 | **99.895** | 98.08 | 8858 | 79746 |
| Dataset 2 | Falcon | 98 | 11083 | 6.994 | 94.58 | 3263 | 47211 |
| Dataset 2 | PBcR | 22 | 246681 | 0.593 | 93.7 | 14823 | 269053 |
| Dataset 2 | ALLPATHS-LG | 2 | **4639001** | **99.938** | **99.99** | 10 | 12 |
| Dataset 2 | SPAdes | 18 | 4488904 | 99.912 | 99.98 | 427 | 110 |
| Dataset 3 | LQS | 3 | **4603990** | **99.998** | **98.49** | 4568 | 65283 |
| Dataset 3 | Falcon | 124 | 13838 | 17.316 | 94.97 | 3206 | 59638 |
| Dataset 3 | PBcR | 1 | 4329903 | 12.825 | 94.03 | 7209 | 262357 |
| Dataset 3 | ALLPATHS-LG | 3 | **4638937** | **99.938** | **99.99** | 6 | 38 |
| Dataset 3 | SPAdes | 19 | 4474608 | 99.88 | 99.99 | 425 | 108 |
| Dataset 4 | LQS | 8 | **4622531** | **99.938** | **99.08** | 2256 | 40118 |
| Dataset 4 | Falcon | 13 | 4538244 | 99.938 | 97.66 | 3710 | 104165 |
| Dataset 4 | PBcR | 3 | 3615068 | 99.553 | 97.39 | 2394 | 117397 |
| Dataset 4 | ALLPATHS-LG | 1 | **4638952** | **99.938** | **99.99** | 8 | 20 |
| Dataset 4 | SPAdes | 20 | 4475770 | 99.88 | 99.99 | 399 | 73 |
| Dataset 5 | LQS | 5 | 4006324 | **99.991** | 99.43 | 1435 | 25106 |
| Dataset 5 | Falcon | 1 | 4580230 | 99.655 | 98.84 | 2589 | 50662 |
| Dataset 5 | PBcR | 1 | 4596475 | 99.914 | 98.99 | 1136 | 45542 |
| Dataset 5 | ALLPATHS-LG | 1 | **4638958** | **99.938** | **99.99** | **3** | **5** |
| Dataset 5 | SPAdes | 16 | **4648863** | **99.918** | **99.99** | 420 | 53 |

Looking at the table, it can be concluded that hybrid assembly pipelines achieve better results than non-hybrid ones. However, this is mostly because Illumina reads provide additional coverage of the genome. ALLPATHS-LG has better results than SPAdes on all datasets, with SPAdes almost catching up on Dataset 5.

None of the non-hybrid assembly pipelines managed to complete the genome at 20x coverage. LQS pipeline produced the best assembly – it managed to cover almost the whole genome, albeit using 8 separate contigs. 30x seems to be sufficient for LQS pipeline to get very good results and for PBcR to cover most of the genome, however with only one contig which is notably shorter than the reference genome. On the other hand, Falcon seems to require at least 40x to produce one big contig covering most of the reference genome. One surprising result that can be seen in the table is a noticeable drop in assembly quality for LQS pipeline on Dataset 5. While it managed to cover a greater part of the reference than any other pipeline on any dataset, with the assembly consists of 5 contigs, the largest of which is just over 4Mbp.

Furthermore, in the “big contig” analysis where only one largest contig of length ≥ 4Mbp (a representative of the E. coli chromosome) was selected and evaluated using Dnadiff. This analysis gave a good estimate on the quality of the assembly from the aspects of chromosome completeness and breakage. Concretely, PBcR had the largest number of breakpoints on Datasets 3 and 4 (≥ 200; for Datasets 1 and 2 its assembly did not produce a “big contig”), while on Dataset 5 LQS had the largest number of breakpoints (88; see **Suppl. Table 2**).

Since the results for all three non-hybrid assembly tools show notable variation in assembly quality across datasets (**Table 1**), we further investigated the differences between their pipelines. As described in the Background section, there are two major differences: (I) LQS and PBcR both employ WGS (Celera) as their middle-layer assembler while Falcon implements its own string graph layout module, and (II) each of these pipelines implements its own error-correction module. Taking into account that both Celera and Falcon utilize an overlap-graph based layout step, we suspected that (II) may have played a more significant role on the assembly contiguity. The error-correction process is performed very early in each pipeline, and the quality of corrected reads can directly influence any downstream analysis. For this purpose, we analysed the error rates in raw reads from Dataset3 as well as the error-corrected reads generated by Nanocorrect, PBcR and Falcon’s error-correction modules (**Suppl. Fig. 1**). For analysis, all reads were aligned to the E. coli K-12 reference (NC_000913.3) using GraphMap (parameters “-a anchorgotoh”). The results show that each method produces significantly different error profile of the corrected reads. The raw dataset (coverage 28.78x) contained a mixture of ~3% insertions, ~4% deletions and ~9% mismatches. While the insertion errors were mostly eliminated by all error-correctors, PBcR and Falcon exhibited higher amounts of deletion errors in their output. Nanocorrect produced the best results, reducing both deletion and mismatch rates to 1%, while still maintaining a large coverage of the output error-corrected reads (25.85x).

To assess the influence of the difference (I), we used the error-corrected reads generated by Nanocorrect as the input data for Falcon for every dataset. We noticed that this procedure increased both the contiguity of Falcons assembly and the average identity on all datasets (**Suppl. Table 3**). Increase in coverage provided a consistent increase of the quality of assembly in terms of the largest contig length, the average identity and the number of variants. Although the draft assemblies produced by the LQS pipeline exhibited a reduction in the size of the largest contig on Dataset5, these assemblies also resulted in a lower number of variants (SNPs and indels) compared to the Nanocorrect+Falcon combination.

### 4.2 Resource usage

To estimate efficiency of each assembly pipeline, the CPU time and the maximum memory usage were measured. All assembly pipelines consumed fewer than 20GB of memory. The memory consumption was mostly stable and is not shown. To conclude, for assembling bacterial genomes, available memory should not be an issue.

**Table 2** shows how CPU usage changes with dataset coverage for each assembly pipeline. SPAdes proved to be the fastest of the tested assemblers, while LQS was the most time consuming.

**Table 2.** CPU time usage in hours

|  | Coverage |  |  |  |
| --- | --- | --- | --- | --- |
| Assembler | 20 | 30 | 40 | 50 |
| LQS | 1085.9 | 2538.6 | 4437.7 | 8142.1 |
| ALLPATHS-LG | 13.2 | 26.0 | 44.9 | 144.5 |
| PBcR | 6.2 | 13.7 | 14.1 | 19.3 |
| Falcon | 3.1 | 6.4 | 19.7 | 13.8 |
| SPAdes | 0.9 | 1.0 | 1.1 | 1.2 |

### 4.3 Hybrid pipeline comparison

Hybrid and non-hybrid assembly pipelines are not directly comparable (except by comparing absolutely best cases) because hybrid pipelines have an advantage in greater coverage supplied by Illumina reads. **Table 3** gives a more detailed comparison between two hybrid assemblers ALLPATHS-LG and SPAdes. Besides running both pipelines on Dataset 0 (pared-end and mate-pair reads) together with each nanopore dataset, SPAdes was also tested using only Illumina paired-end reads (without mate-pair reads).

**Table 3.** Comparing ALLPATHS-LG and SPAdes results

| **Dataset** | **Assembler** | **# ctg** | **N50** | **Genome fraction (%)** | **Avg. Identity 1-to-1** | **Total SNPs** | **Total Indels** |
| --- | --- | --- | --- | --- | --- | --- | --- |
| Dataset 0 | ALLPATHS-LG | 3 | **4626283** | **99.219** | **99.99** | 59 | 73 |
| Dataset 0 | SPAdes PE and MP | 106 | 1105151 | 99.089 | 99.98 | 231 | 78 |
| Dataset 2 | ALLPATHS-LG | 2 | **4639001** | **99.938** | **99.99** | **10** | **12** |
| Dataset 2 | SPAdes PE only | 20 | 4470699 | 99.908 | 99.99 | 430 | 93 |
| Dataset 2 | SPAdes PE and MP | 18 | 4488904 | 99.912 | 99.98 | 427 | 110 |
| Dataset 3 | ALLPATHS-LG | 3 | **4638937** | **99.938** | **99.99** | **6** | **38** |
| Dataset 3 | SPAdes PE only | 19 | 4474624 | 99.908 | 99.99 | 418 | 92 |
| Dataset 3 | SPAdes PE and MP | 19 | 4474608 | 99.88 | 99.99 | 425 | 108 |
| Dataset 4 | ALLPATHS-LG | 1 | **4638952** | **99.938** | **99.99** | **8** | **20** |
| Dataset 4 | SPAdes PE only | 20 | 4475777 | 99.908 | 99.99 | 401 | 66 |
| Dataset 4 | SPAdes PE and MP | 20 | 4475770 | 99.88 | 99.99 | 399 | 73 |
| Dataset 5 | ALLPATHS-LG | 1 | **4638958** | **99.938** | **99.99** | **3** | **5** |
| Dataset 5 | SPAdes PE only | 18 | **4648869** | **99.918** | **99.99** | 421 | 47 |
| Dataset 5 | SPAdes PE and MP | 16 | **4648863** | **99.918** | **99.99** | 420 | 53 |

The table shows that ALLPATHS-LG produces better results than SPAdes on all datasets, from Dataset 0 without nanopore data, for which SPAdes is not able to produce one sufficiently large contig, to Dataset 5 on which the difference is miniscule and apparent only in the number of SNPs and indels.

It is interesting to notice that while ALLPATHS-LG requires both a paired-end and a mate-pair library to run, SPAdes seems not to be able to leverage mate-pair reads at all. Results using paired-end Illumina library without mate-pairs seems to be equal to or even slightly better than with a mate-pair library, for all nanopore datasets. This means that in a situation where expensive-to-produce mate-pairs reads are unavailable, SPAdes might be a good choice for a de novo assembler.

## 5 Conclusion

In our study we compared several hybrid and non-hybrid de novo assembly tools and assessed their ability to work with nanopore data. Each examined tool proved capable of assembling a whole bacterial genome under the right conditions. The choice of the best assembly tool will heavily depend upon the characteristics of the datasets. ALLPATHS-LG showed overall best results, but it requires both paired-end and mate-pair short reads. In case only paired-end reads are available, SPAdes might be the better choice. Of the non-hybrid assembly tools, on some datasets LQS pipeline came close to or even surpassed hybrid tools. However, extremely high CPU time used by the Nanocorrect might make it prohibitively slow on larger genomes and larger datasets, in which case Falcon or PBcR could be used instead. Additionally, the good results obtained from hybrid assembly tools were largely due to higher coverage provided by Illumina reads.

We can expect that with further development of nanopore technology (and other long read sequencing technologies) read quality will increase and the technology will become more accessible and more affordable. This will make *de novo* assembly using nanopore reads faster, more precise and applicable to lager genomes.

## 6 Funding

This work has been supported in part by Croatian Science Foundation under the project UIP-11-2013-7353 Algorithms for Genome Sequence Analysis.

